# Widespread impact of immunoglobulin V gene allelic polymorphisms on antibody reactivity

**DOI:** 10.1101/2023.06.06.543969

**Authors:** Meng Yuan, Ziqi Feng, Huibin Lv, Natalie So, Ivana R. Shen, Timothy J.C. Tan, Qi Wen Teo, Wenhao O. Ouyang, Logan Talmage, Ian A. Wilson, Nicholas C. Wu

## Abstract

The ability of human immune system to generate antibodies to any given antigen can be strongly influenced by immunoglobulin V gene (IGV) allelic polymorphisms. However, previous studies have provided only a limited number of examples. Therefore, the prevalence of this phenomenon has been unclear. By analyzing >1,000 publicly available antibody-antigen structures, we show that many IGV allelic polymorphisms in antibody paratopes are determinants for antibody binding activity. Biolayer interferometry experiment further demonstrates that paratope allelic mutations on both heavy and light chain often abolish antibody binding. We also illustrate the importance of minor IGV allelic variants with low frequency in several broadly neutralizing antibodies to SARS-CoV-2 and influenza virus. Overall, this study not only highlights the pervasive impact of IGV allelic polymorphisms on antibody binding, but also provides mechanistic insights into the variability of antibody repertoires across individuals, which in turn have important implications for vaccine development and antibody discovery.

## INTRODUCTION

Human antibodies, which are produced by B cells and composed of two chains (heavy and light), are central to the immune response against pathogen infection. To be able to recognize many different pathogens, the human antibody repertoire has an enormous diversity with up to 10^15^ unique antibody clonotypes [1]. This diversity is generated mainly by V(D)J recombination, which is a somatic recombination process that randomly assembles different germline gene segments, known as variable (V), diversity (D), and joining (J) genes, into the variable region of the antibody molecule. In addition, many V gene segments are known to have multiple alleles with amino acid differences, which further increase the diversity of human antibody repertoire at the population level.

Allelic polymorphisms of immunoglobulin V (IGV) genes are enriched in the complementarity-determining regions (CDRs) [2], which usually form the antibody paratopes (i.e. regions that involve in binding to antigens). Several studies have reported the importance of IGV allelic polymorphisms in antibody binding. For example, allelic polymorphisms at IGHV1-69 residues 50 (G/R) and 54 (F/L) (n.b. Kabat numbering used throughout for all antibody residues) can influence antibody binding to severe acute respiratory syndrome coronavirus 2 (SARS-CoV-2) [3]. Allelic polymorphisms at IGHV1-69 residues 50 (G/R) can also impact antibody binding to *Staphylococcus aureus* (*S. aureus*) [4]. Other examples include IGHV3-33 residue 52 (W/S) in antibodies to *Plasmodium falciparum* (*P. falciparum*) [5] and IGHV2-5 residue 54 (D/N) in antibodies to SARS-CoV-2 [6] and human immunodeficiency virus (HIV) [7]. Therefore, it is apparent that IGV allelic polymorphisms can affect antibody binding. However, it remains unclear whether the impact of IGV allelic polymorphisms on antibody binding is prevalent, since previous studies typically characterized a single antibody-antigen pair at a time and the number of such studies is rather limited. Furthermore, previous studies of IGV allelic polymorphisms mainly focused on the heavy chain [3–10], whereas those within the light chain remain largely unexplored.

Nevertheless, IGV allelic polymorphisms are shown to have important public health relevance. For instance, IGHV1-2 allelic usage correlates with the response rate to an HIV vaccine candidate in a phase 1 clinical trial (NCT03547245) [8, 9, 11], due to the influence of its allelic polymorphisms at residue 50 (W/R) on antibody binding to HIV [8, 12, 13]. A previous study has also shown that IGHV1-69 allelic usage correlates with the broadly neutralizing antibody response to influenza virus in a vaccine cohort [14]. This observation has also been attributed to the differential autoreactive propensities of dfferent IGHV1-69 alleles [10] as well as a potentially minor effect of its allelic polymorphisms at residue 54 (L/F) on antibody binding to the conserved stem domain of influenza hemagglutinin (HA) [15, 16]. Similarly, allele usages of IGHV3-66 and IGHV4-61 were associated with Kawasaki disease [17] and rheumatic heart disease [18], respectively, although the underlying mechanisms are unknown. As a result, investigating the impact of IGV allelic polymorphisms can provide critical insights into vaccine development and autoimmune diseases [19].

In this study, we systematically investigate the effect of IGV allelic polymorphisms on antibody binding by analyzing 1,398 publicly available antibody-antigen complex structures. IGV allelic polymorphisms could be identified in the antibody paratope of 39% (544/1,398) complex structures. Computational analysis of protein mutational stability predicted that 73% of paratope allelic mutations (i.e. mutating a paratope residue to alternative allelic variants) would weaken antibody binding and 19% were strongly disruptive. The wide impact of IGV allelic polymorphisms on antibody binding was further validated using biolayer interferometry (BLI). Our results also illustrated the importance of light chain IGV allelic polymorphisms. In addition, we identified several minor IGV allelic variants with low frequency that are essential for the binding activity of broadly neutralizing antibodies to SARS-CoV-2 and influenza virus.

## RESULTS

### Predicting IGV allelic mutations that weaken antibody binding activity

To systematically analyze the impact of IGV allelic polymorphisms on antibody binding activity, we leveraged the collection of antibody-antigen complex structures available in the Structural Antibody Database (SAbDab, http://opig.stats.ox.ac.uk/webapps/sabdab) [20]. Among 1,398 antibody-antigen complex structures that we analyzed, 544 contained at least one paratope residue with IGV allelic polymorphism (**see Methods**). Most antigens in these 544 structures, which had a median resolution of 2.6 Å (range = 1.2 Å to 7.3 Å, **Figure S1A**), were from viruses, although a considerable number were from human and *Plasmodium* (**Figure S1B**). Subsequently, we computationally predict the impact of paratope allelic polymorphisms in these 544 structures on antibody binding activity (**Figure 1A**). For example, there are three allelic variants at residue 50 of IGHV4-4, namely Arg, Glu, and Tyr. If an antibody is encoded by IGHV4-4 and had V_H_ Arg50 in the paratope, we would predict the effects of allelic mutations V_H_ R50E and V_H_ R50Y on its binding activity. In this study, we predicted the effects of 1,150 paratope allelic mutations across 544 structures on antibody binding activity (**see Methods and Table S1**).

**Figure 1.**
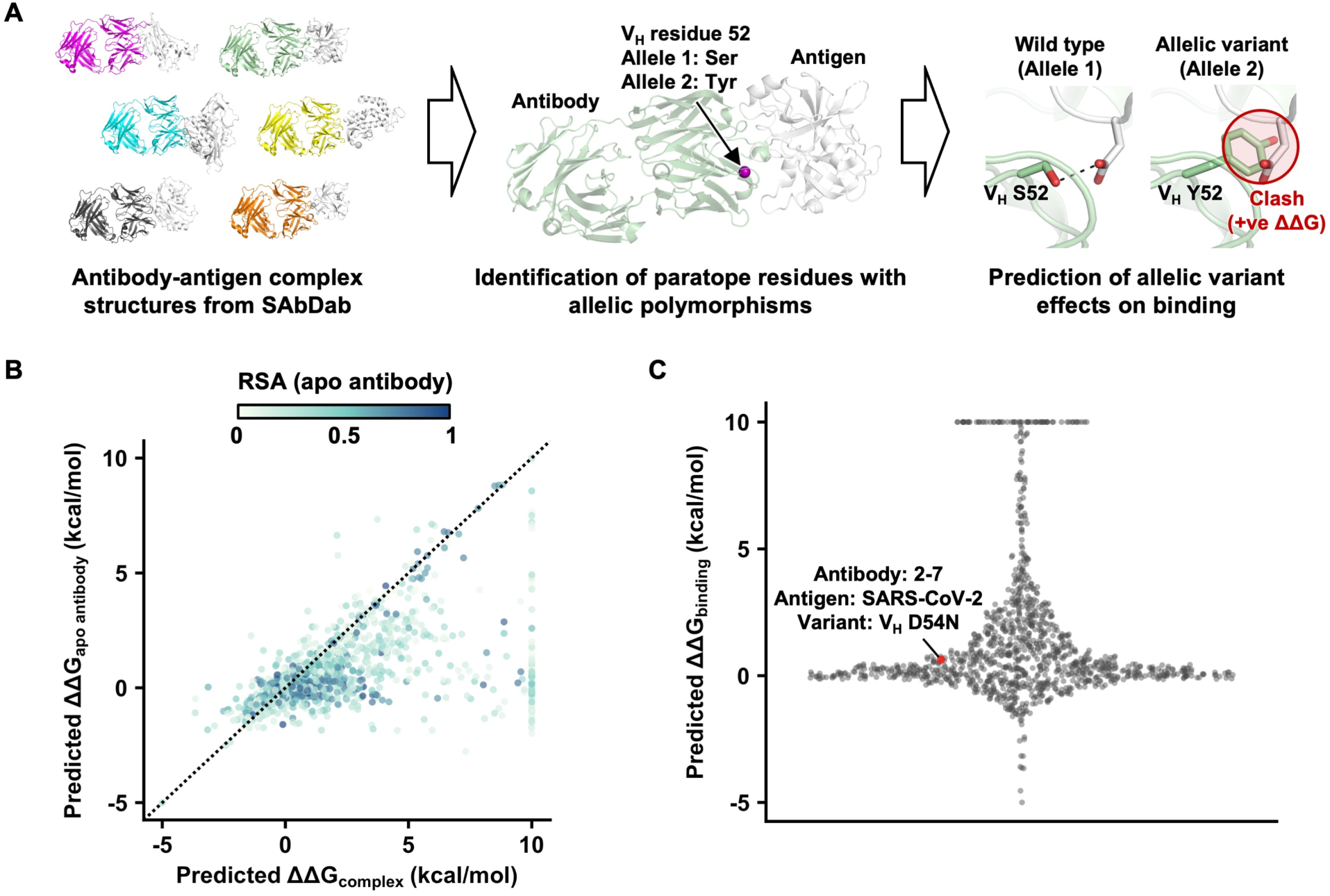
Predicting the effects of IGV allelic mutations on antibody binding activity. **(A)** Schematic of the analysis workflow (**see Methods**). Briefly, human antibody-antigen complex structures were downloaded as PDB files from the Structural Antibody Database (SAbDab, http://opig.stats.ox.ac.uk/webapps/sabdab) [20]. Among these antibodies, paratope residues with allelic polymorphisms were identified. The effects of allelic mutations on antibody binding activity were then predicted using FoldX [21]. **(B)** The relationship between predicted ΔΔG_apo antibody_ and predicted ΔΔG_complex_ is shown. Each data point is colored by the relative solvent accessibility (RSA) in the apo antibody. Residues that are fully solvent exposed have an RSA of 1, whereas those that are fully buried have an RSA of 0. **(C)** The distribution of predicted ΔΔG_binding_ of 1,150 paratope allelic mutations of 544 antibody-antigen complex structures is shown. Predicted ΔΔG_binding_ was computed by predicted ΔΔG_complex_ – predicted ΔΔG_apo antibody_. Mutations with predicted ΔΔG > 10 kcal/mol are shown as 10 kcal/mol. Mutations with predicted ΔΔG < -5 kcal/mol are shown as -5 kcal/mol.

A paratope allelic mutation could impact the stability of the antibody-antigen complex structure (ΔΔG_complex_) by altering the stability of the antibody (ΔΔG_apo antibody_) as well as the antibody-antigen binding energy (ΔΔG_binding_). Therefore, to predict ΔΔG_binding_, we would need to first predict both ΔΔG_complex_ and ΔΔG_apo antibody_ (**Figure S1C-D**). Here, we used FoldX [21] to predict the ΔΔG_complex_ and ΔΔG_apo antibody_ of each of the 1,150 paratope allelic mutations. The predicted ΔΔG_binding_ was then calculated by subtracting predicted ΔΔG_apo antibody_ from predicted ΔΔG_complex_. Here, ΔΔG > 0 kcal/mol indicated destabilization. Many paratope allelic mutations had a higher predicted ΔΔG_complex_ than predicted ΔΔG_apo antibody_ (i.e. predicted ΔΔG_binding_ > 0 kcal/mol, **Figure 1B-C**). Of note, predicted ΔΔG_binding_ had minimal correlation with the resolution of the structures (rank correlation = -0.16, **Figure S1E**), indicating that quality of the structures did not systematically bias the estimation of ΔΔG_binding_.

### Paratope allelic polymorphisms often affect antibody binding activity

A previous benchmarking study has shown that FoldX has a 70% accuracy of classifying whether a mutation is stabilizing (ΔΔG < 0 kcal/mol) or destabilizing (ΔΔG > 0 kcal/mol) [22]. Here, we considered a predicted ΔΔG_binding_ of >2.5 kcal/mol as strong disruption of antibody-antigen binding, with 0 kcal/mol to 2.5 kcal/mol as mild disruption [23, 24]. Among the 1,150 paratope allelic mutations from 544 structures, 19% (217 mutations from 177 structures) were predicted to be strongly disruptive and 54% (620 mutations from 398 structures) to be mildly disruptive. Although FoldX is known to be more accurate at predicting destabilizing mutations than stabilizing mutations [23], we acknowledge that FoldX may misclassify some mutations that improve or have no effect on binding (i.e. ΔΔG_binding_ ≤ 0 kcal/mol) as disruptive (i.e. ΔΔG > 0 kcal/mol). At the same time, the impact of certain paratope allelic mutations on antibody binding activity may be underestimated by FoldX. For example, while the binding dissociation constant (K_D_) of IGHV2-5 antibody 2-7 to the receptor-binding domain of SARS-CoV-2 spike was shown to be >100-fold weaker with the IGHV2-5 allelic mutation D54N [6], its predicted ΔΔG_binding_ was only 0.63 kcal/mol, corresponding to only around 3-fold change in K_D_ (**Figure 1C**).

Next, we examined whether the impact of allelic polymorphisms on antibody binding activity could be observed in different IGV genes. Within the 544 antibody-antigen complex structures that were analyzed, paratope allelic mutations were identified in 43 IGV genes. Among these 43 IGV genes, 88% (38/43) and 60% (26/43) had at least one paratope allelic mutation with a predicted ΔΔG_binding_ > 0 kcal/mol and a predicted ΔΔG_binding_ > 2.5 kcal/mol, respectively (**Figure 2A**). These IGV genes spanned different IGHV, IGKV, and IGLV families. Similarly, we also observed that paratope allelic mutations affected antibody binding to many different antigens (**Figure 2B**). For example, 18% (139/776) and 11% (5/45) of paratope allelic mutations in antibodies to viral and bacterial antigens, respectively, had a predicted ΔΔG_binding_ > 2.5 kcal/mol. We also observed that 25% (58/232) of paratope allelic mutations in antibodies to human proteins had a predicted ΔΔG_binding_ > 2.5 kcal/mol. These antibodies to human proteins include several FDA-approved therapeutic antibodies, namely avelumab (PDB 5GRJ) [25], dupilumab (PDB 6WGL) [26], tralokinumab (PDB 5L6Y) [27], atezolizumab (PDB 5XXY) [28], dostarlimab (PDB 7WSL) [29], and daratumumab (PDB 7DHA) [30]. Overall, these observations suggest that IGV allelic polymorphisms play a critical role in determining the binding activity of antibodies with different germline usages and specificities.

**Figure 2.**
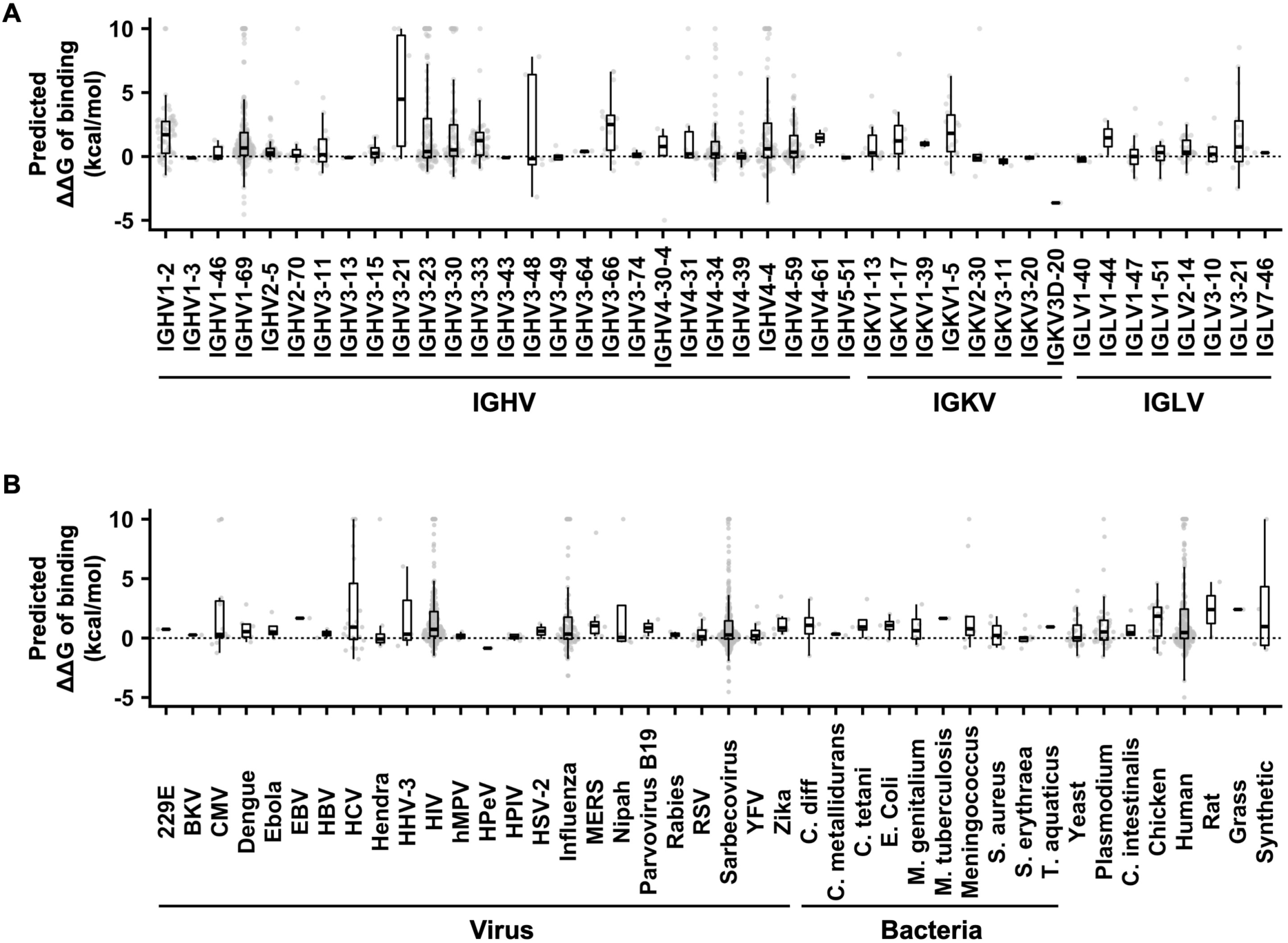
IGV allelic polymorphisms influence the binding activity of diverse antibodies. The distributions of predicted ΔΔG_binding_ of paratope allelic mutations in **(A)** different IGV genes and **(B)** antibodies to different antigens are shown. One data point represents one paratope allelic mutation. For the boxplot, the middle horizontal line represents the median. The lower and upper hinges represent the first and third quartiles, respectively. The upper whisker extends to the highest data point within a 1.5× inter-quartile range (IQR) of the third quartile, whereas the lower whisker extends to the lowest data point within a 1.5× IQR of the first quartile. Mutations with predicted ΔΔG > 10 kcal/mol are shown as 10 kcal/mol. Mutations with predicted ΔΔG < -5 kcal/mol are shown as -5 kcal/mol.

### Experimentally validating the importance of paratope allelic polymorphisms

To further confirm the prevalent impact of IGV allelic polymorphisms in antibody paratopes, we experimentally determined the effect of 14 paratope allelic mutations on antibody binding affinity. All these 14 paratope allelic mutations had a predicted ΔΔG_binding_ > 0 (**Table S2**), 11 of which had a predicted ΔΔG_binding_ > 2.5. They were selected among antibodies that bind to the antigens of five medically important pathogens, namely SARS-CoV-2 spike, HCV E2 envelope glycoprotein, HIV Env, influenza HA, and *P. falciparum* circumsporozoite protein (CSP). In addition, these 14 paratope allelic mutations were on eight different IGV genes and had different biophysical properties, such as charge reversion (IGHV4-4 R50E), decrease in side-chain volume (IGHV3-33 W52S), and increase in side-chain volume (IGHV3-30 V50F). Our BLI experiment showed that among these 14 paratope allelic mutations, 10 abolished the antibody binding activity, whereas the remaining four weakened the binding affinity by at least 5-fold (**Figure 3A, Figure S2, and Figure S4**). These findings substantiate that paratope allelic polymorphisms have a prevalent impact on the binding activity of diverse antibodies.

**Figure 3.**
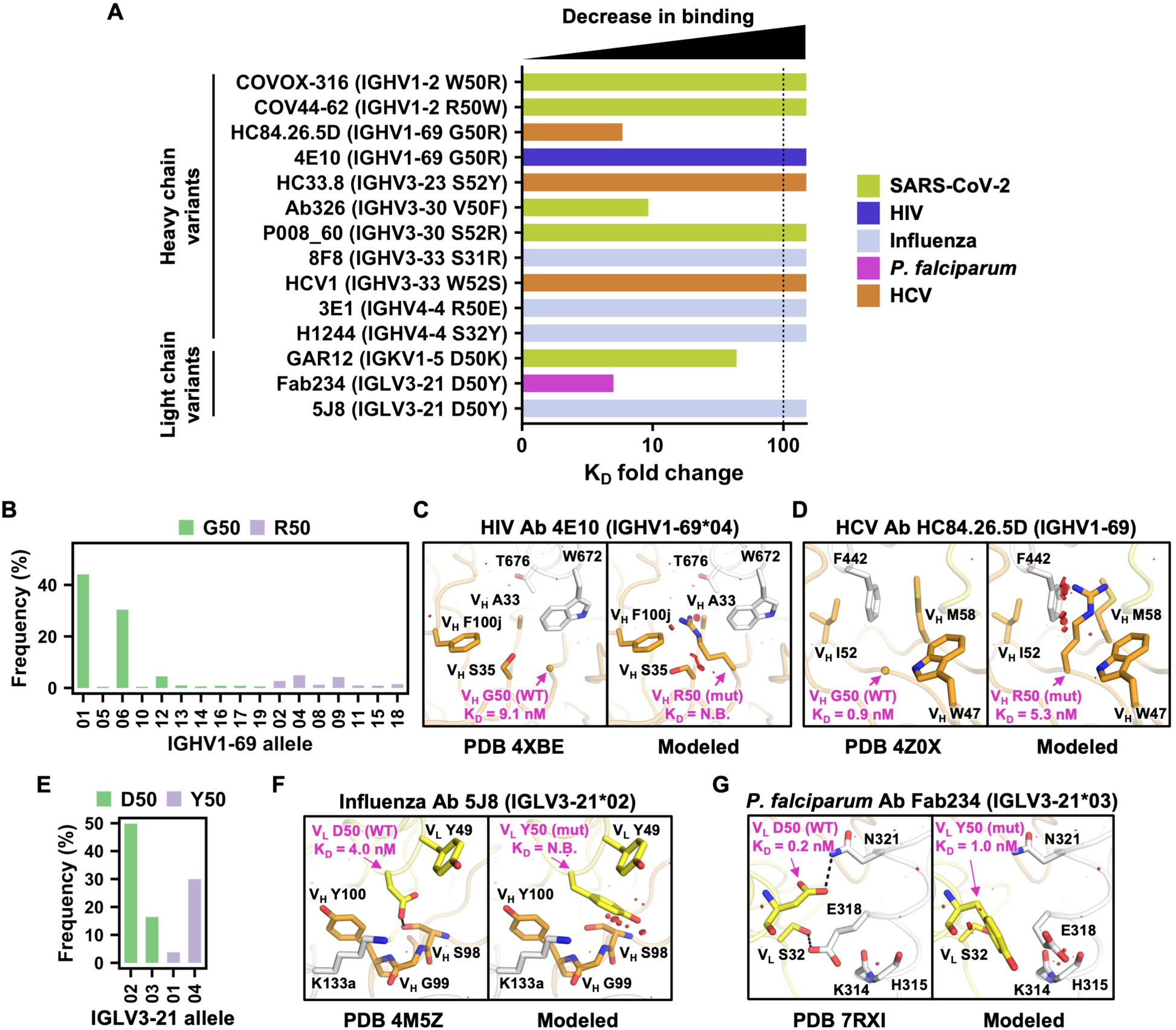
A given allelic polymorphism can impact the binding activity of antibodies to different antigens. **(A)** The impact of the indicated paratope allelic mutations on the antibody binding affinity of the corresponding antibodies is quantified by the fold-change in the binding dissociation constant (K_D_). A higher fold-change indicates worse binding. Paratope allelic mutations that abolished antibody binding are shown with a fold-change of >100. Antibody specificity is color-coded. **(B, E)** Allele usages of **(B)** IGHV1-69 antibodies, and **(E)** IGLV3-21 antibodies are shown. Bar color represents the amino-acid identity at the indicated residue position. **(C-D and F-G)** Structural effects of paratope allelic mutations **(C)** V_H_ G50R of antibody 4E10 in complex with HIV Env membrane-proximal external region (PDB 4XBE) [52], **(D)** V_H_ G50R of antibody HC84.26.5D in complex with HCV glycoprotein E2 (PDB 4Z0X) [35], **(F)** V_L_ D50Y of antibody 5J8 in complex with influenza H1N1 A/California/07/2009 HA1 subunit (PDB 4M5Z) [53], and **(G)** V_L_ D50Y of Fab234 in complex with C-terminal αTSR domain of *P. falciparum* circumsporozoite protein (PDB 7RXI) [54] are modeled by FoldX [21]. Left panel: previously determined experimental structures of antibody-antigen complexes. Right panel: models of allelic mutations. Allelic polymorphic residues and mutations are labelled in magenta. K_D_ values of wild type and allelic mutants to antigens were measured by biolayer Interferometry (BLI) and are indicated. N.B. represents no binding. Antibody heavy and light chains are shown in orange and yellow, respectively. Antigens are shown in white. Red disks indicate significant van der Waals overlap (distance < 2.8 Å), hence representing a steric clash. H-bonds are represented by black dashed lines. Kabat numbering is applied to all antibodies. Cα atoms of glycines are represented by spheres. The IGV gene and allele usage for each antibody are indicated. Of note, the allele usage for HC84.26.5D cannot be assigned unambiguously.

### A given allelic polymorphism can impact diverse antibodies

IGHV1-69 is frequently utilized in the antibody response against microbial pathogens [31, 32]. Among all of the paratope allelic mutations in IGHV1-69, G50R has an exceptional high predicted ΔΔG_binding_ across multiple antibodies (**Figure S3**). IGHV1-69 encodes either Gly or Arg at residue 50, depending on the allele. Among 1,266 IGHV1-69 antibodies from GenBank [33], 84% were encoded by alleles with Gly50 (**Figure 3B**). Previous studies have shown that mutation V_H_ G50R would abolish the binding activity of IGHV1-69 antibodies to *S. aureus* [4]. Our BLI experiment further demonstrated that V_H_ G50R abolished binding of IGHV1-69 antibody 4E10 to the membrane-proximal external region (MPER) of HIV Env and reduced binding of another IGHV1-69 antibody, HC84.26.5D, to HCV E2 by around 6-fold (**Figure 3A and Figure S2**). Of note, 4E10 represents a multidonor class of IGHV1-69/IGKV3-20 broadly neutralizing antibodies to HIV [34]. Structural modeling showed that V_H_ G50R would introduce substantial steric clash between 4E10 and HIV Env (**Figure 3C**). A similar observation was made for V_H_ G50R in antibody HC84.26.5D (PDB 4Z0X) [35] (**Figure 3D**). A recent preprint has reported that V_H_ G50R can also abolish the binding activity of another IGHV1-69 antibody to HCV E2 [36], although its epitope differs from that of HC84.26.5D. These findings demonstrate that multiple IGHV1-69 antibodies in the literature with different specificities have a strong preference towards alleles that encode Gly50 rather than Arg50.

Another example was IGLV3-21 residue 50, which has either Asp or Tyr, depending on the allele. Among 347 IGLV3-21 antibodies from GenBank [33], 66% were encoded by alleles with Asp50 (**Figure 3E**). Our BLI experiment showed that V_L_ D50Y abolished the binding activity of IGLV3-21 antibody 5J8 to influenza H1N1 A/California/07/2009 HA and weakened the binding affinity of another IGLV3-21 antibody, Fab234, to *P. falciparum* CSP by around 5-fold (**Figure 3A and Figure S2**). Structural modeling indicated that V_L_ D50Y would remove a H-bond and introduce steric clashes between 5J8 and influenza HA (**Figure 3F**), and similarly remove a H-bond between Fab234 and *P. falciparum* CSP (**Figure 3G**). These observations not only substantiate that a given allelic polymorphism can impact the binding activities of diverse antibodies, but also demonstrate that such a phenomenon also arises in light chain.

### Importance of minor allelic variants in broadly neutralizing antibodies

The binding activities of three out of 14 antibodies in our BLI experiment (**Figure 3A**) were attributed to minor IGV allelic variants (i.e. allelic variant frequency < 25%, **Table S2**). These three antibodies, namely GAR12, COV44-62, and 3E1, are all broadly neutralizing antibodies. GAR12 targets the receptor-binding domain of SARS-CoV-2 spike and was previously shown to neutralize all tested variants of concern, including Omicron BA1, BA2, and BA5 [37]. The light chain of GAR12 is encoded by allele 1 of IGKV1-5, which has a minor allelic variant Asp50. Among 744 IGKV1-5 antibodies from GenBank [33], only 16% were encoded by alleles with Asp50 (allele 1 or 2), whereas the remaining 84% were encoded by allele 3, which had Lys50 (**Figure 4A**). Our BLI experiment showed that the binding of GAR12 to the receptor-binding domain of SARS-CoV-2 spike was reduced by 44-fold when V_L_ Asp50 was mutated to the major allelic variant V_L_ Lys50. Structural modeling indicated that V_L_ D50K would disrupt an extensive electrostatic interaction network with Arg346 of the SARS-CoV-2 spike RBD and introduce unfavorable electrostatic interactions (**Figure 4B**).

**Figure 4.**
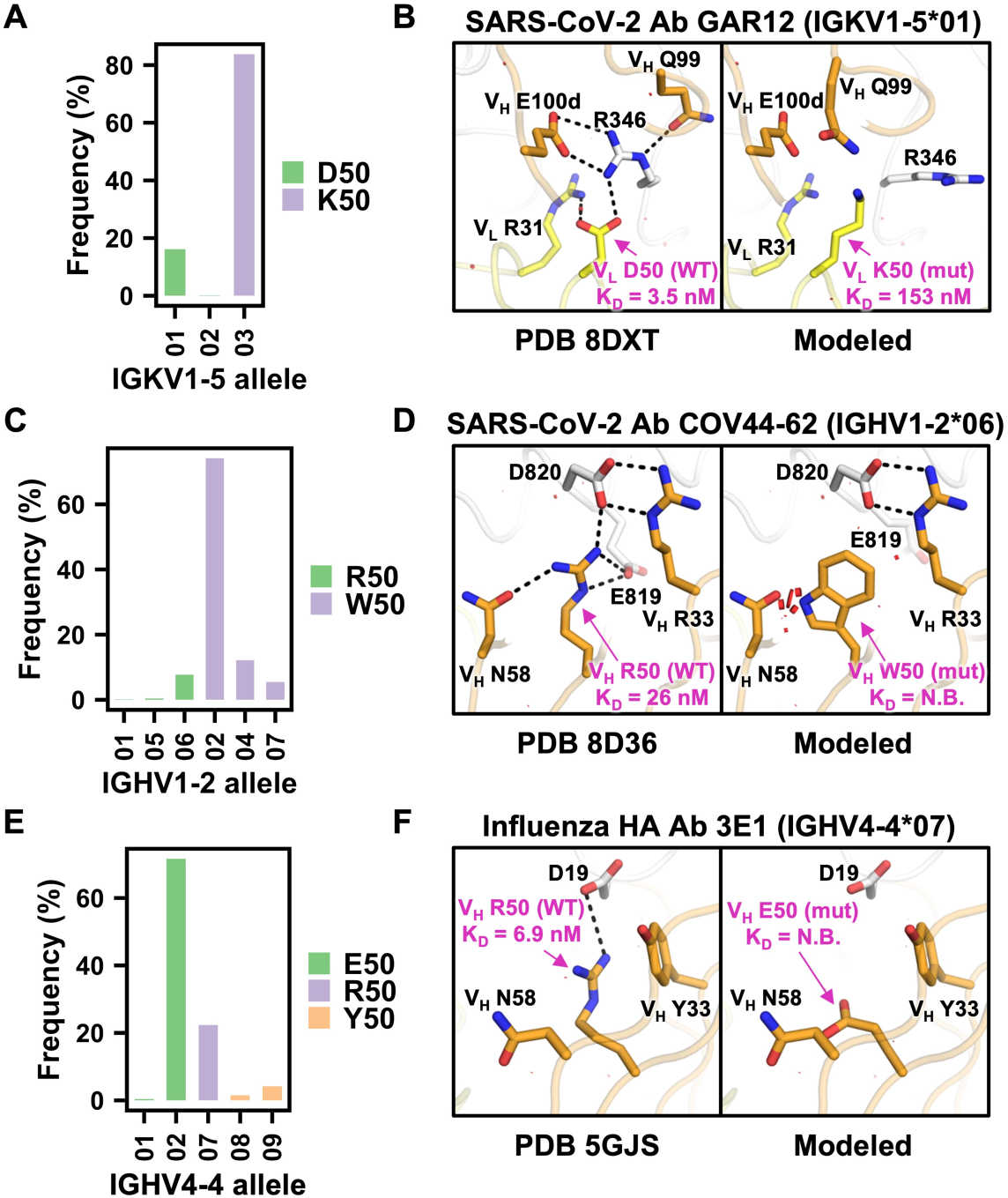
Minor IGV allelic variants are important for the binding activity of diverse antibodies. **(A, C, E)** Allele usage of **(A)** IGKV1-5 antibodies, **(C)** IGHV1-2 antibodies, and **(E)** IGHV4-4 antibodies is shown. Bar color represents the amino-acid identity at the indicated residue position. **(B, D, F)** Structural effects of paratope allelic mutations **(B)** V_L_ D50K of antibody GAR12 in complex with the receptor-binding domain of SARS-CoV-2 spike (PDB 8DXT) [37], **(D)** V_H_ R50W of antibody COV44-62 in complex with the fusion peptide of SARS-CoV-2 spike (PDB 8D36) [38], and **(F)** V_H_ R50E of antibody 3E1 in complex with influenza H1N1 A/California/04/2009 HA (PDB 5GJS) [39] are modeled by FoldX [21]. Structure visualization has the same style and format as Figure 3, with H-bonds and salt bridges represented by black dashed lines.

COV44-62 targets the highly conserved fusion peptide of coronavirus spike and was previously shown to be neutralize coronavirus strains from different genera [38]. The heavy chain of COV44-62 is encoded by allele 6 of IGHV1-2, which has a minor allelic variant Arg50. Among 788 IGHV1-2 antibodies from GenBank [33], only 8% were encoded by alleles with Arg50, whereas the remaining 92% were encoded by alleles with Trp50 (**Figure 4C**). Our BLI experiment showed that binding of COV44-62 to the fusion peptide of SARS-CoV-2 spike was abolished by mutating V_H_ Arg50 to the major allelic variant V_H_ Trp50 (**Figure 3A and Figure S2**). Structural modeling indicated that V_H_ R50W would remove multiple salt bridges between COV44-62 and SARS-CoV-2 fusion peptide as well as introduce steric clashes at the binding interface (**Figure 4D**).

3E1 targets the conserved stem domain of influenza HA and was previously shown to neutralize influenza A H1 and H5 subtypes [39]. The heavy chain of 3E1 is encoded by IGHV4-4, which has a minor allelic variant Arg50. Among 264 IGHV4-4 antibodies from GenBank [33], 6%, 22%, and 72% were encoded by alleles with Tyr50, Arg50, and Glu50, respectively (**Figure 4E**). Our BLI experiment showed that the binding of 3E1 to influenza H1N1 A/California/07/2009 HA was abolished by mutating V_H_ Arg50 to the major allelic variant V_H_ Glu50 (**Figure 3A and Figure S2**). Structural modeling showed that V_H_ R50E would eliminate a salt bridge between 3E1 and influenza HA (**Figure 4F**). Overall, these results demonstrate the contribution of minor IGV allelic variants to broadly neutralizing antibody responses against different antigens.

## DISCUSSION

Previous studies have provided limited examples of how IGV allelic polymorphisms can affect the binding activity of antibodies of interest [3–8]. By analyzing more than a thousand publicly available antibody-antigen complex structures, our study here shows that the impact of IGV allelic polymorphisms on antibody binding activity is highly prevalent. Consistent with these findings, a recent preprint reports that IGV allelic polymorphisms are linked to the variability of antibody repertoires across individuals [40]. These observations indicate that IGV allelic polymorphisms can greatly influence antibody responses to different antigens. Consequently, the potency and breadth of antibody response that is elicited by infection or vaccination may be associated with IGV allele usage. This phenomenon is indeed observed in a Phase I clinical trial of an HIV vaccine candidate [8, 9] and likely generalizable. Given that certain broadly neutralizing antibodies rely on minor IGV allelic variants, IGV allelic polymorphisms in the human population are an important consideration in the pursuit of more universal vaccines.

The need, if any, for maintaining multiple alleles of a given IGV gene in the human population has been previously regarded as an evolutionary mystery [2]. One possibility is that different alleles of a given IGV gene have different antigen-binding preferences, which would in turn lead to heterozygote advantage. An example is residue 50 of IGHV1-69, which encodes either Gly or Arg, depending on alleles. V_H_ G50R abolished the binding activity of IGHV1-69 antibodies to *S. aureus*, as shown in a previous study [4], as well as HIV Env and HCV E2, as shown in our work here. Nevertheless, a recent study has shown that V_H_ Arg50 is essential for IGHV1-69 antibodies to interact with the receptor-binding domain of SARS-CoV-2 spike [3]. As a result, mounting an optimal IGHV1-69 antibody response against *S. aureus*, HIV, HCV, and SARS-CoV-2 would require both Gly50 and Arg50 alleles of IGHV1-69. In other words, individuals with either Gly50 or Arg50 in all copies of IGHV1-69 gene may have difficulties generating effective IGHV1-69 antibody response against certain pathogens. Another example is residue 50 of IGHV1-2, which encodes either Arg or Trp, depending on alleles. Our work here showed that V_H_ Arg50 is essential for the binding activity of COV44-62, which is a IGHV1-2 broadly neutralizing coronavirus antibody [38]. In contrast, V_H_ Trp50 is essential for the binding activity of IGHV1-2 broadly neutralizing HIV antibodies [8, 12, 13]. Based on our ΔΔG_binding_ prediction results, similarly observations can be made for allelic polymorphisms in other germline genes (**Figure S3**). Nevertheless, future experimental studies will be needed to fully dissect the evolutionary causes and consequences of allelic polymorphisms in different IGV genes.

In this study, we also identified several FDA-approved therapeutic antibodies where paratope allelic mutations were predicted to strongly disrupt binding to human proteins, including cancer therapeutic targets. These antibodies were identified by phage display screening of human antibody libraries (e.g. avelumab [41], tralokinumab [42], and atezolizumab [43]) and immunization of humanized mice (e.g. dupilumab [44] and daratumumab [45]), which are two common methods for antibody discovery. Both methods require cloning of antibody repertoires from human donors. Based on our results, it is likely that the probability of success of antibody discovery through phage display screening and humanized mice depends not only on the antigens, but also IGV allelic polymorphisms of the human donors present in the phage-displayed antibody libraries or in the humanized mice. Future antibody discovery may benefit from having multiple human donors with diverse IGV alleles. Given that the market size of monoclonal antibodies continues to grow [46], understanding the impact of IGV allelic polymorphisms on antibody discovery will have important public health implications.

While this study indicates that many IGV allelic polymorphisms can influence antibody binding activity, our analysis only focused on those in the paratope. Previous studies have demonstrated that non-paratope mutations can also affect antibody binding activity [47–49]. Therefore, some non-paratope IGV allele polymorphisms may possibly do the same. Besides, the diversity of human IGV alleles is likely higher than what is currently known. Over the past decade, numerous IGV alleles have been discovered thanks to the advances in computational methods for germline gene inference and long-read third-generation sequencing technologies [19]. The ongoing efforts in understanding the diversity of IGV genes across human populations and ethnicities, including geographic diversity, will likely reveal many more novel IGV alleles [50]. Of note, the impact of IGV allelic polymorphisms on antibody binding are unlikely to be limited to human IGV genes, since a similar observation has been recently reported in IGHV3-73 of rhesus macaque to SARS-CoV-2 [51]. Together, it is likely that IGV allelic polymorphisms, as well as their impact on antibody binding activity, are more widespread than indicated by our and previous studies [19].

## ACKNOWLEDGEMENTS

This work was supported by the National Institutes of Health (NIH) R01 AI167910 (N.C.W.), DP2 AT011966 (N.C.W.), the Bill and Melinda Gates Foundation INV-004923 (I.A.W.), UM1 AI144462 (I.A.W.), Department of Health and Human Services under contract number: 75N93021C00015 (I.A.W., N.C.W.), and the Searle Scholars Program (N.C.W.). We thank Jeanne Matteson and Beverly Ellis for contribution to mammalian cell culture, Wenli Yu, Xueyong Zhu, Re’em Moskovitz, Tossapol Pholcharee, and T.K. Yen Nguyen for assistance in protein production. We are grateful for the SARS-CoV-2 spike (HexaPro) plasmid from Jason McLellan from The University of Texas at Austin.

## AUTHOR CONTRIBUTIONS

M.Y. and N.C.W. conceived and designed the study. N.S. and N.C.W. analyzed the antibody structure database and performed mutational stability analysis. M.Y. and N.C.W. performed the structural analysis. M.Y., Z.F., H.L., I.R.S., T.J.C.T., Q.W.T., W.O.O., and L.T. expressed and purified the antibodies and antigens. M.Y. and Z.F. performed the biolayer interferometry experiment. N.C.W. and I.A.W. provided resources and support. M.Y. and N.C.W. wrote the paper and all authors reviewed and/or edited the paper.

## METHODS

### Identification of paratope residues with allelic polymorphisms

A total of 3,240 human antibody-antigen complex structures were downloaded as PDB files from the Structural Antibody Database (SAbDab, http://opig.stats.ox.ac.uk/webapps/sabdab) [20]. Next, PDB files with more than one antibody were filtered out, leaving 1,416 complex structures. We further discarded the following PDB files due to formatting issues: 7T1W, 7T1X, 6TUL, 6SS4, 6SS5, 7DWT, 7DWU, 6SS2, 6ZJG, 7T0W, 6YXM, 6TKF, 6TKE, 6TKD, 6TKC, 3J6U, 7R8U, and 6YXL, leaving 1,398 complex structures. For each PDB file, an apo antibody structure was generated by removing the antigen from the PDB file. Relative solvent accessibility (RSA) for each antibody residue, either in apo form or in complex with antigen, was computed by DSSP [55]. Residues with a higher RSA value in the apo antibody structure than the complex structure (i.e. RSA_apo antibody_ – RSA_complex_ > 0) were defined as paratope residues. Germline sequences for human IGHV and IGK(L)V genes were downloaded from the IMGT database (https://www.imgt.org/) [56]. Each position of the antibody sequences in the PDB files and the germline IGV gene sequences was numbered according to Kabat numbering using ANARCI [57].

The germline IGV genes of each antibody were identified using PyIR [58], a wrapper for the IgBLAST [59]. For each paratope residue, its allelic variants across different alleles of the corresponding germline IGV gene were compared. Residues with two or more allelic variants were defined as residues with allelic polymorphisms. Paratope residues with no allelic polymorphism were excluded from downstream analysis. In addition, if the amino-acid identity of a given paratope residue did not match any of its germline amino-acid variants, such a paratope residue was also discarded. If a paratope residue has two alternative allelic variants, two allelic mutations at that given paratope residue were investigated. In summary, a total of 1,150 paratope allelic mutations across 544 antibody-antigen complex structures were identified.

### Predicting the ΔΔG of binding for allelic mutations

The ΔΔG for each paratope allelic mutations was predicted using FoldX [21]. For each paratope allelic mutation of a given antibody, two ΔΔG values were predicted, one for the apo antibody structure (ΔΔG_apo antibody_) and the other for the antibody-antigen complex structure (ΔΔG_complex_). Apo antibody structures were generated by extracting the antibody coordinates from the PDB files. Predicted ΔΔG of antibody-antigen binding (ΔΔG_binding_) was computed as:

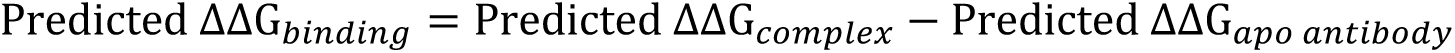

### Germline IGV gene and allele assignment of antibodies from GenBank

The germline IGV genes and alleles of 12,487 antibodies from GenBank (www.ncbi.nlm.nih.gov/genbank) [33] were identified using PyIR [58].

### Expression and purification of Fabs

The heavy and light chains of Fabs were cloned into phCMV3 vector. PCR-based mutagenesis was performed to generate the allelic mutations. The plasmids were transiently co-transfected into ExpiCHO cells at a ratio of 2:1 (heavy chain to light chain) using ExpiFectamine CHO Reagent (Thermo Fisher Scientific) according to the manufacturer’s instructions. The supernatant was collected at 7 days post-transfection. The Fab was purified with a CaptureSelect CH1-XL Pre-packed Column (Thermo Fisher Scientific), followed by a buffer exchange to Dulbecco’s Phosphate Buffered Saline (PBS, pH 7.4).

### Expression and purification of antigens

SARS-CoV-2 receptor-binding domain was expressed in High Five cells and purified with Ni-NTA resin followed by size exclusion as described previously [60]. SARS-CoV-2 whole spike protein HexaPro was a gift from Jason McLellan (Addgene plasmid # 154754). SARS-CoV-2 HexaPro was expressed in Expi293F cells and purified with Ni-NTA resin followed by size exclusion as described previously [61]. The influenza hemagglutinin (HA) proteins from H1N1 A/California/07/2009, H1N1 A/Beijing/262/1995, and H2N2 A/Japan/305/1957 were expressed in High Five cells, purified with Ni-NTA resin followed by size exclusion, and biotinylated as described previously [62]. The HCV (isolate H77) E2 domain was expressed in HEK293S cells and purified as described previously [63]. The biotinylated SARS-CoV-2 fusion peptide (N’-biotin-DPSKPSKRSFIEDLLFNKVT-C’) and His-tagged HIV Env MPER peptide (N’-NWFDITNWLWYIKSGGSHHHHHHHH-C’) were chemically synthesized by GenScript.

### Biolayer Interferometry (BLI) binding assays

Binding assays were performed by biolayer interferometry (BLI) using an Octet Red instrument (FortéBio). 20 μg/ml of antibodies, antigens, or peptides in 1x kinetics buffer (1x PBS, pH 7.4, 0.01% BSA and 0.002% Tween 20) were loaded onto different types of sensors, and then incubated with 33 nM, 100 nM, and 300 nM of binders. All antibodies were in Fab format. Specifically, biotinylated SARS-CoV-2 fusion peptide was loaded onto Streptavidin (SA) sensors and incubated with COV44-62 (WT or V_H_ R50W). Biotinylated αTSR domain of *P. falciparum* was loaded onto SA sensors and incubated with Fab234 (WT or V_L_ D50Y). Biotinylated influenza HA from H1N1 A/California/07/2009 was loaded onto SA sensors and incubated with 3E1 (WT or V_H_ R50E), or 5J8 (WT or V_L_ D50). Biotinylated influenza HA from H1N1 A/Beijing/262/1995 was loaded onto SA sensors and incubated with H1244 (WT or V_H_ S32Y). Biotinylated virus HA from H2N2 A/Japan/305/1957 was loaded onto SA sensors and incubated with 8F8 (WT or V_H_ S31R). His_6_-tagged SARS-CoV-2 receptor-binding domain was loaded onto Ni-NTA sensors and incubated with GAR12 (WT or V_L_ D50K), Ab326 (WT or V_H_ V50F), or COVOX-316 (WT or V_H_ W50R). His-tagged HIV Env MPER peptide was loaded onto Ni-NTA sensors and incubated with 4E10 (WT or V_H_ G50R). HC33.8 (WT or V_H_ S52Y), HC84.26.5D (WT or V_H_ G50R), or HCV1 (WT or V_H_ W52S) was loaded onto Anti-Human Fab-CH1 2nd Generation (FAB2G) sensors and incubated with HCV E2. P008_60 (WT or V_H_ S52R) was loaded onto FAB2G sensors and incubated with SARS-CoV-2 spike protein [61]. The assay consisted of five steps: 1) baseline; 2) loading; 3) baseline; 4) association; and 5) dissociation. For estimating the K_D_ values, a 1:1 binding model was used.

### Code availability

Custom python scripts for all analyses have been deposited to: https://github.com/nicwulab/Ab_allele_polymorphism.

