## Supplementary Information for "Widespread impact of immunoglobulin V gene allelic polymorphisms on antibody reactivity"

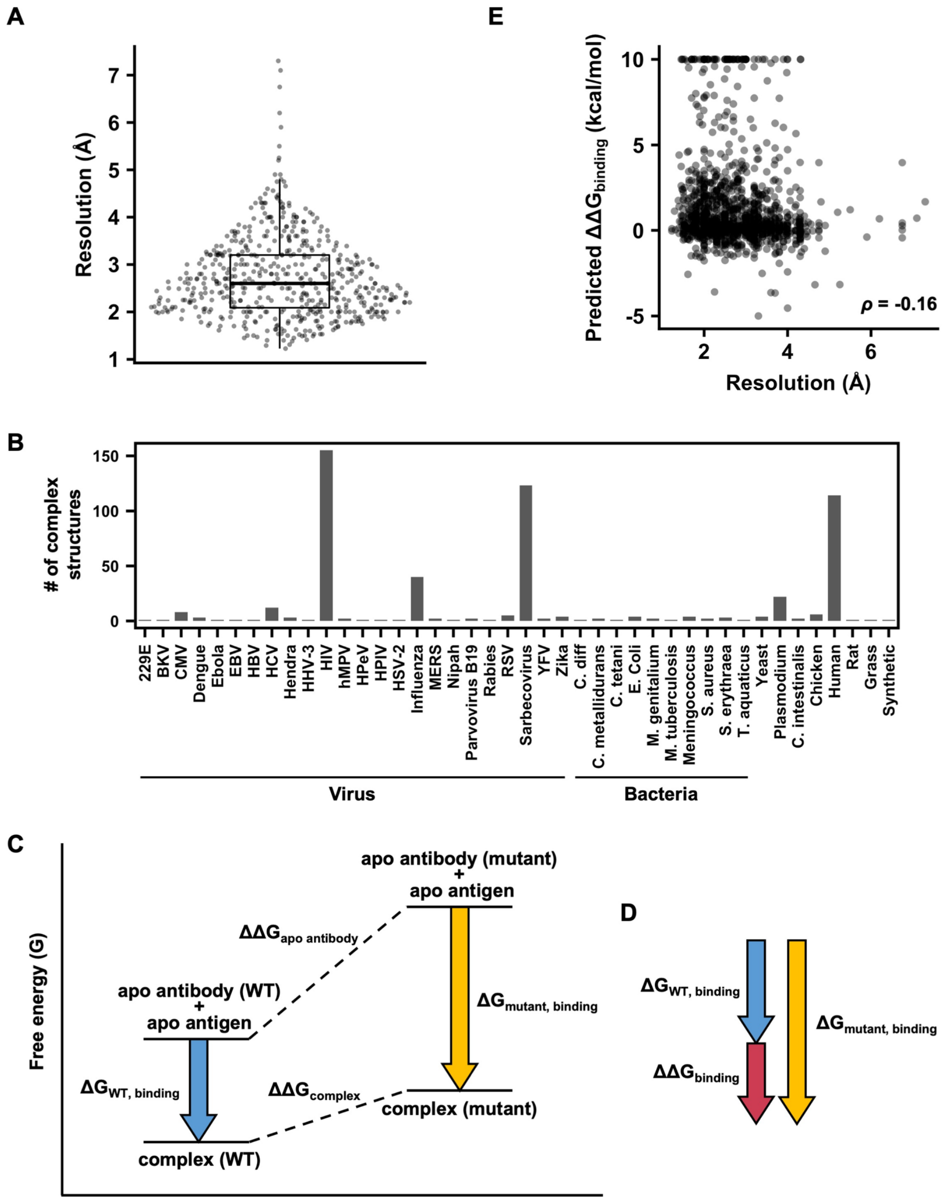


**Figure S1. Overview of the dataset and ΔΔG calculation. (A)** Resolutions (in Å) of the 544 antibody-antigen complex structures are shown. **(B)** Paratope residues with allelic polymorphisms were identified in 544 antibody-antigen complex structures. The species of the antigen among these 544 antibody-antigen complex structures are plotted, with the occurrence frequency shown on the y-axis. **(C)** A free energy diagram of binding between antibody and antigen is shown. Upon mutation of the antibody, the free energy (G) of both the complex and the apo antibody may change, which can be quantified as $\Delta\Delta G_{complex}$ and $\Delta\Delta G_{apo\ antibody}$, respectively. Blue arrow indicates the $\Delta G$ of binding of the wild-type (WT) antibody ($\Delta G_{WT,\ binding}$), whereas the yellow arrow indicates the $\Delta G$ of binding of the mutant antibody ($\Delta G_{mutant,\ binding}$). **(D)** The difference between $\Delta G_{WT,\ binding}$ and $\Delta G_{mutant,\ binding}$ is represented by $\Delta\Delta G_{binding}$, which can be quantified as $\Delta\Delta G_{complex} - \Delta\Delta G_{apo\ antibody}$. That is, $\Delta\Delta G_{binding} = \Delta G_{mutant,\ binding} - \Delta G_{WT,\ binding} = \Delta\Delta G_{complex} - \Delta\Delta G_{apo\ antibody}$. In this example here, the mutation strengthens the binding since it destabilizes the apo antibody to a greater extent than the complex. **(E)** The relationship between resolution and $\Delta\Delta G_{binding}$ of the 1,150 paratope allelic mutations is shown. The Spearman's rank correlation coefficient ($\rho$) is indicated. Mutations with predicted $\Delta\Delta G > 10$ kcal/mol are shown as 10 kcal/mol. Mutations with predicted $\Delta\Delta G < -5$ kcal/mol are shown as -5 kcal/mol. (**A and E**) One data point represents one paratope allelic mutation.

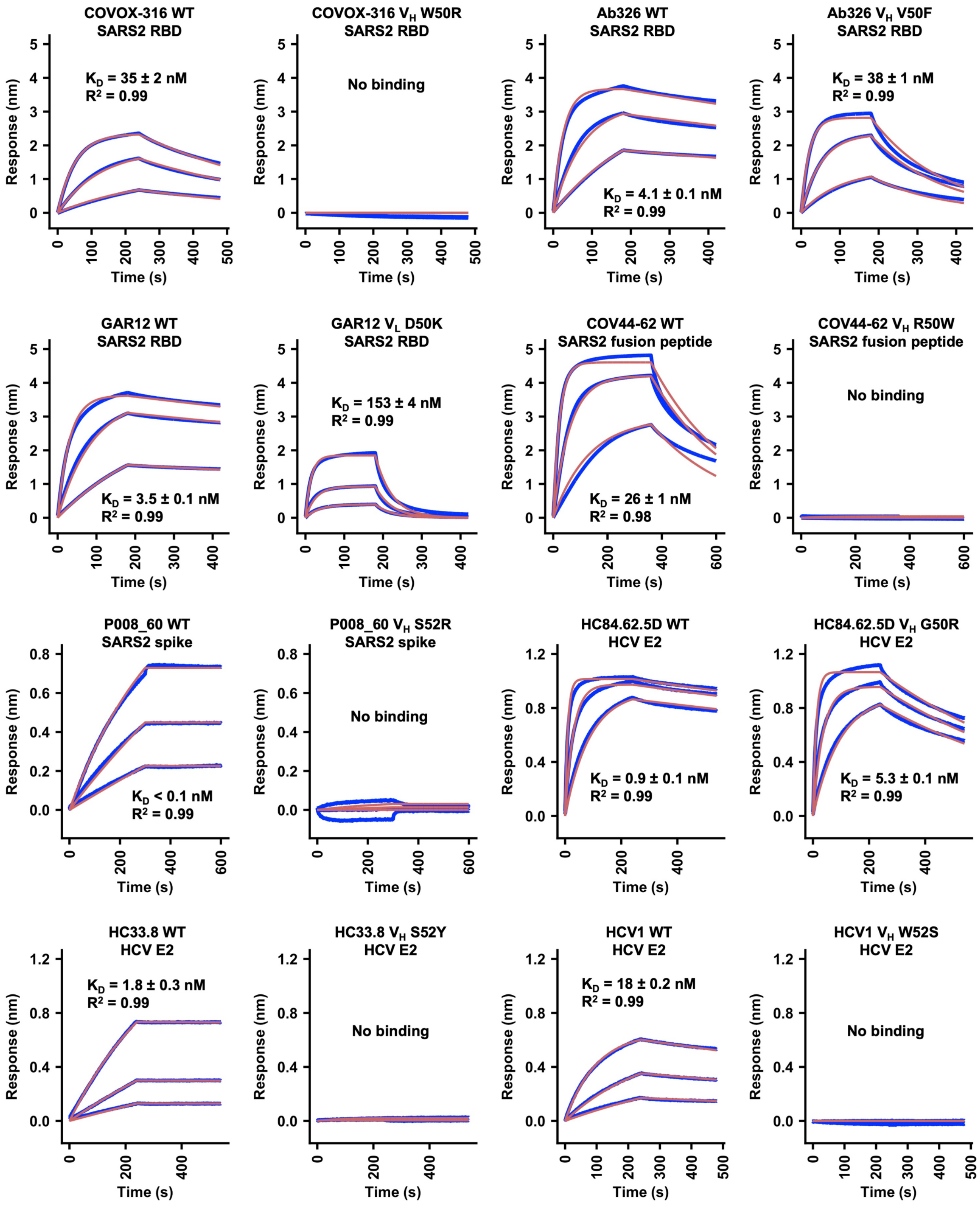
COVOX-316 WT
SARS2 RBD
$K_D$ = 35 ± 2 nM
$R^2$ = 0.99
COVOX-316 $V_H$ W50R
SARS2 RBD
No binding
Ab326 WT
SARS2 RBD
$K_D$ = 4.1 ± 0.1 nM
$R^2$ = 0.99
Ab326 $V_H$ V50F
SARS2 RBD
$K_D$ = 38 ± 1 nM
$R^2$ = 0.99
GAR12 WT
SARS2 RBD
$K_D$ = 3.5 ± 0.1 nM
$R^2$ = 0.99
GAR12 $V_L$ D50K
SARS2 RBD
$K_D$ = 153 ± 4 nM
$R^2$ = 0.99
COV44-62 WT
SARS2 fusion peptide
$K_D$ = 26 ± 1 nM
$R^2$ = 0.98
COV44-62 $V_H$ R50W
SARS2 fusion peptide
No binding
P008_60 WT
SARS2 spike
$K_D$ < 0.1 nM
$R^2$ = 0.99
P008_60 $V_H$ S52R
SARS2 spike
No binding
HC84.62.5D WT
HCV E2
$K_D$ = 0.9 ± 0.1 nM
$R^2$ = 0.99
HC84.62.5D $V_H$ G50R
HCV E2
$K_D$ = 5.3 ± 0.1 nM
$R^2$ = 0.99
HC33.8 WT
HCV E2
$K_D$ = 1.8 ± 0.3 nM
$R^2$ = 0.99
HC33.8 $V_H$ S52Y
HCV E2
No binding
HCV1 WT
HCV E2
$K_D$ = 18 ± 0.2 nM
$R^2$ = 0.99
HCV1 $V_H$ W52S
HCV E2
No binding
Response (nm)
Time (s)

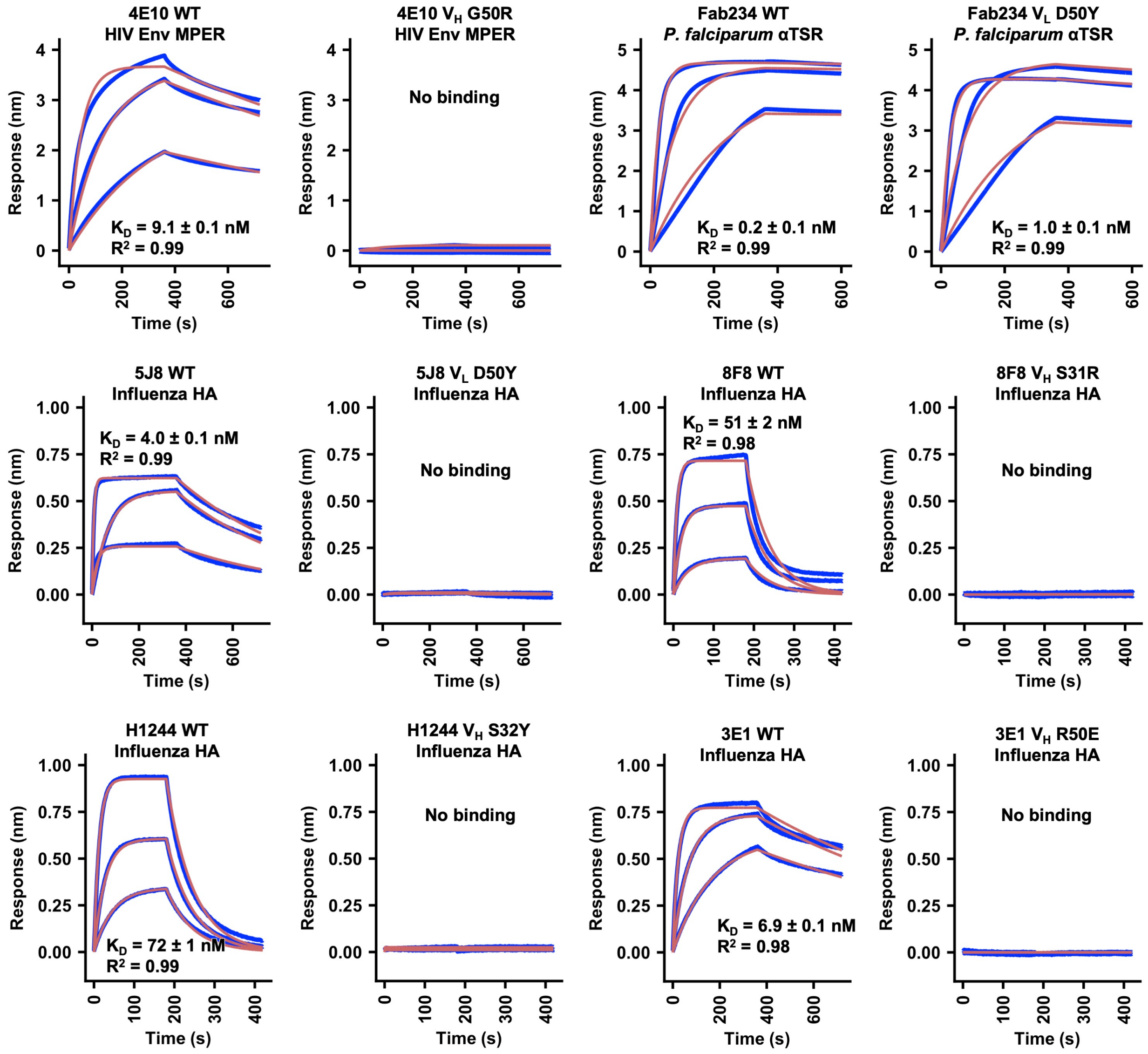


**Figure S2. Sensorgrams for binding of Fabs to recombinantly expressed antigens.** Binding kinetics of different Fabs against their corresponding recombinantly expressed antigens were measured by biolayer interferometry (BLI). Y-axis represents the response. Blue lines represent the response curve and red lines represent a 1:1 binding model. Binding kinetics were measured for three Fab or antigen concentrations. Dissociation constant ($K_D$) and the goodness of model fitting ($R^2$) are indicated.

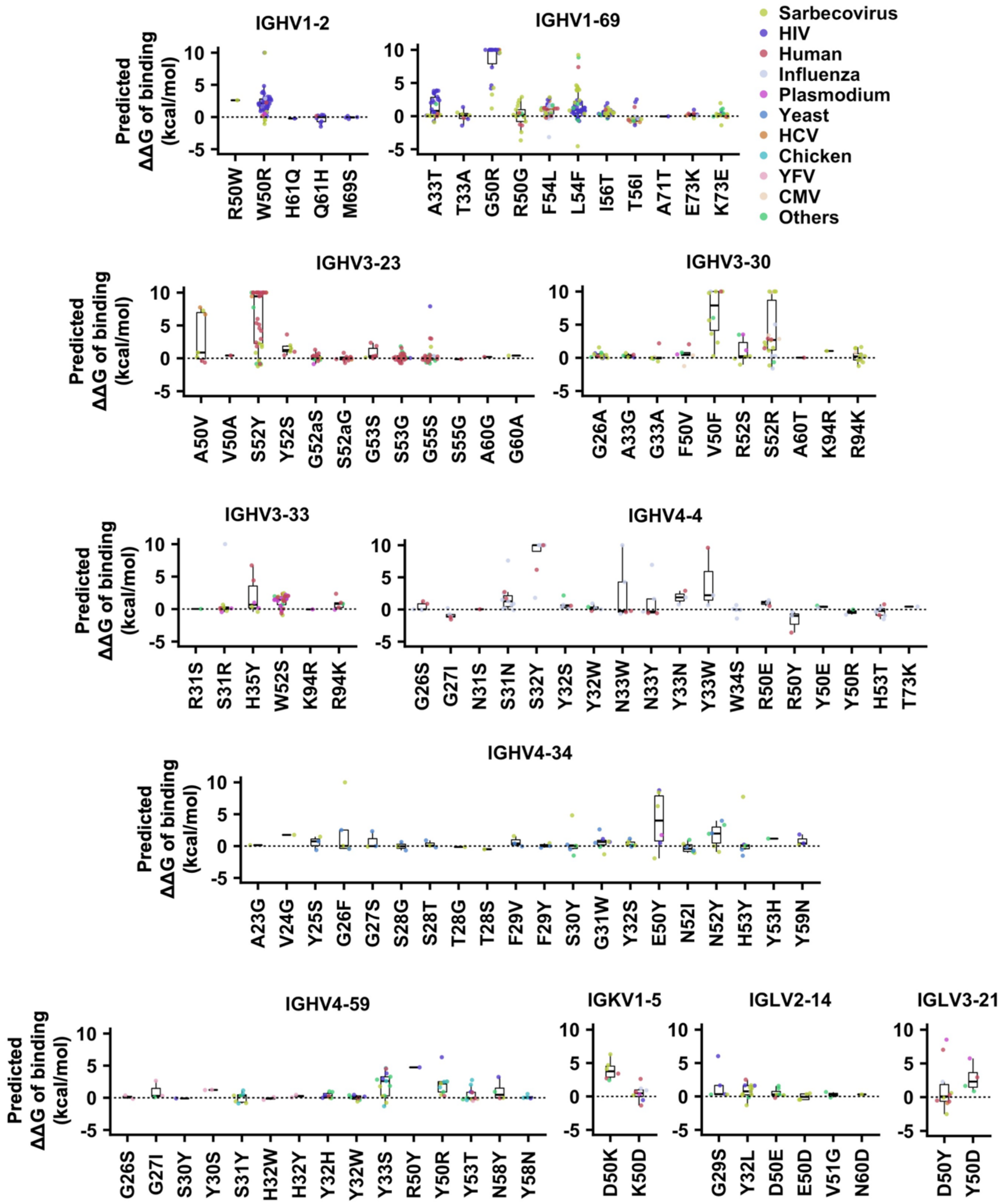


**Figure S3. Distributions of predicted ΔΔG$_{binding}$ of paratope allelic mutations.** The distributions of predicted ΔΔG$_{binding}$ of paratope allelic mutations in antibodies encoded by the indicated IGV genes are shown. Paratope allelic mutations are categorized by their identities and colored by the antigens. One data point represents one paratope allelic mutation. Mutations with predicted ΔΔG > 10 kcal/mol are shown as 10 kcal/mol. Mutations with predicted ΔΔG < -5 kcal/mol are shown as -5 kcal/mol.

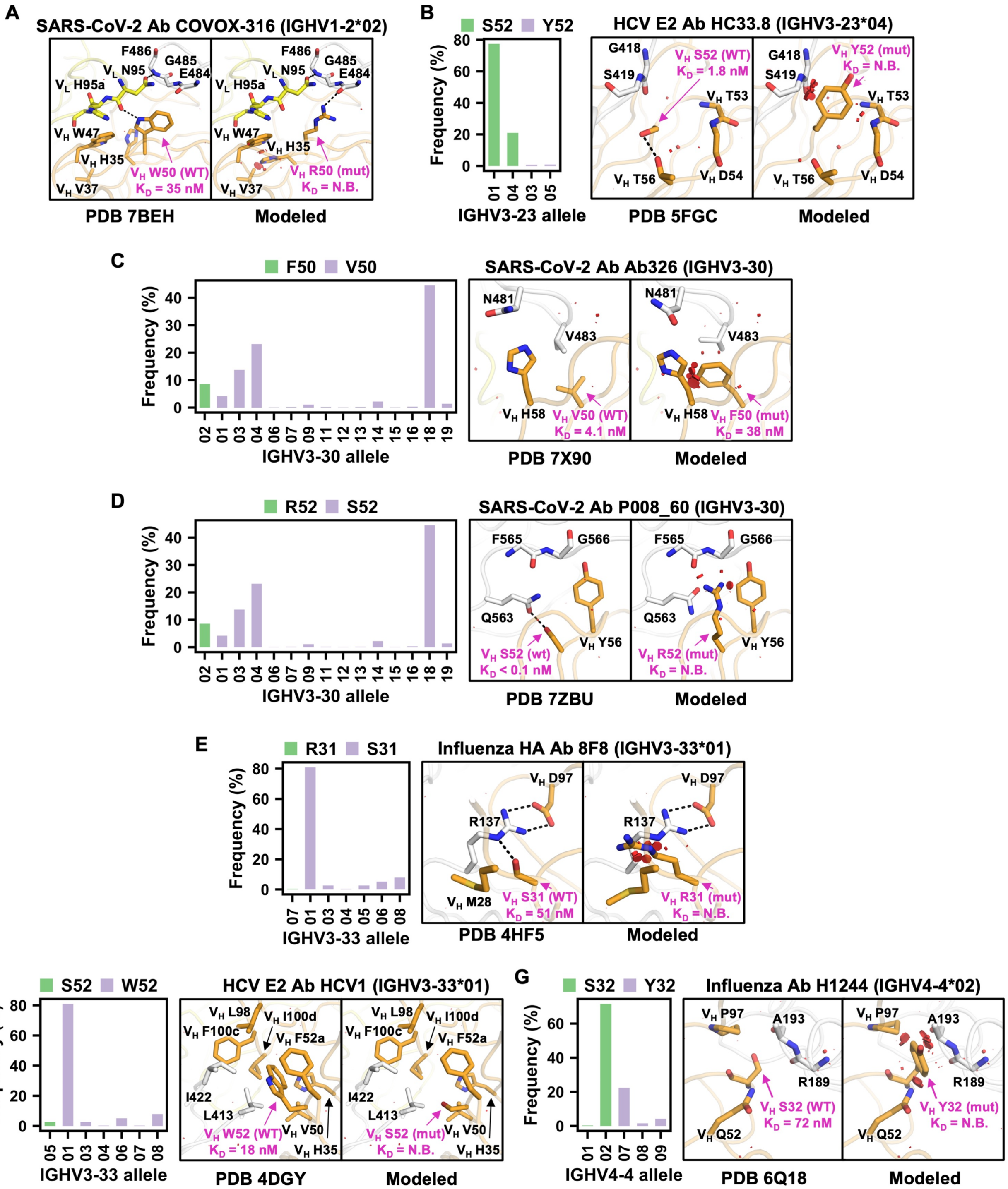


**Figure S4. Structural analysis of selected paratope allelic mutations.** The structural effects of paratope allelic mutations **(A)** $V_H$ W50R of antibody COVOX-316 in complex with the receptor binding domain (RBD) of SARS-CoV-2 spike (PDB 7BEH) [1], **(B)** $V_H$ S52Y of antibody H33.8 in complex with a peptide fragment of HCV E2 (PDB 5FGC) [2], **(C)** $V_H$ V50F of antibody Ab326 in complex with the RBD of SARS-CoV-2 spike (PDB 7X90) [3], **(D)** $V_H$ S52R of antibody P008_60 in complex with SARS-CoV-2 spike monomer (PDB 7ZBU) [4], **(E)** $V_H$ S31R of antibody 8F8 in complex with influenza H2N2 A/Japan/305/1957 hemagglutinin (PDB 4HF5) [5], **(F)** $V_H$ W52S of antibody HCV1 in complex with a peptide fragment of HCV E2 (PDB 4DGY) [6], and **(G)** $V_H$ S32Y of antibody H1244 in complex with influenza H1N1 A/Beijing/262/1995 hemagglutinin head domain (PDB 6Q18) [7], are modeled by FoldX [8]. The IGV gene and allele usage for each antibody are indicated. Of note, the allele usages for Ab326 and P008_60 cannot be assigned unambiguously. Binding dissociation constant ($K_D$) values of wild type and allelic mutant to the antigens were measured by BLI and are indicated. Structure visualization was generated using the same style and format as **Figure 4**. The bar charts on the right in each panel shows the allele usages of different IGV genes. Bar color represents the amino-acid identity at the indicated residue position.

**Table S2. Summary of experimentally validated paratope allelic mutations.**

| PDB | Antibody | Antigen | Allelic mutation | WT freq (%)[a] | Predicted $\Delta\Delta G_{binding}$ (kcal/mol)[b] | WT $K_D$ (nM) | Mutant $K_D$ (nM) |
|---|---|---|---|---|---|---|---|
| 8DXT | GAR12 | SARS-CoV-2 spike | IGKV1-5 D50K | 16.3 | 6.3 | 3.5 | 153 |
| 7BEH | COVOX-316 | SARS-CoV-2 spike | IGHV1-2 W50R | 91.8 | 17.2 | 35 | No binding |
| 7X90 | Ab326 | SARS-CoV-2 spike | IGHV3-30 V50F | 91.5 | 5.7 | 4.1 | 38 |
| 8D36 | COV44-62 | SARS-CoV-2 spike | IGHV1-2 R50W | 8.2 | 2.6 | 26 | No binding |
| 7ZBU | P008_60 | SARS-CoV-2 spike | IGHV3-30 S52R | 91.5 | 11.7 | <0.1 | No binding |
| 4Z0X | HC84.62.5D | HCV E2 | IGHV1-69 G50R | 83.6 | 14.2 | 0.9 | 5.3 |
| 5FGC | HC33.8 | HCV E2 | IGHV3-23 S52Y | 98.4 | 20.6 | 1.8 | No binding |
| 4DGY | HCV1 | HCV E2 | IGHV3-33 W52S | 97.3 | 2.4 | 18 | No binding |
| 4XBE | 4E10 | HIV Env | IGHV1-69 G50R | 83.6 | 12.6 | 9.1 | No binding |
| 5GJS | 3E1 | Influenza HA | IGHV4-4 R50E | 22.3 | 1.4 | 6.9 | No binding |
| 6Q18 | H1244 | Influenza HA | IGHV4-4 S32Y | 72.0 | 22.2 | 72 | No binding |
| 4HF5 | 8F8 | Influenza HA | IGHV3-33 S31R | 99.7 | 17.9 | 51 | No binding |
| 4M5Z | 5J8 | Influenza HA | IGLV3-21 D50Y | 66.3 | 2.3 | 4.0 | No binding |
| 7RXI | Fab234 | *P. falciparum* CSP | IGLV3-21 D50Y | 66.3 | 8.5 | 0.2 | 1.0 |

[a] Occurrence frequency of individual alleles of the indicated germline genes was estimated based on antibody sequences downloaded from GenBank (www.ncbi.nlm.nih.gov/genbank) [9] (**see Methods**). The occurrence frequency of alleles that encode the wild-type (WT) amino acid at the indicated position of the corresponding antibody is listed. For example, antibody GAR12 is encoded by IGKV1-5 with Asp50. Among all the IGKV1-5 antibodies in GenBank, 16.3% were assigned to IGKV1-5 alleles that encode Asp50 in the germline.

[b] $\Delta\Delta G_{binding}$ was predicted using FoldX [8].
